# ExMove: An open-source toolkit for processing and exploring animal tracking data in R

**DOI:** 10.1101/2023.05.16.540922

**Authors:** Liam Patrick Langley, Stephen Lang, Luke Ozsanlav-Harris, Alice Trevail

## Abstract

1. Ongoing technological advances have led to a rapid increase in the number, type, and scope of animal tracking studies. In response, many software tools have been developed to analyse animal movement data. These tools generally focus on movement modelling, but the steps required to clean raw data files from different tracking devices have been largely ignored. Such pre-processing steps are often time-consuming and involve a steep learning curve but are crucial for the creation of high quality, standardised, and shareable data. Moreover, decisions made at this early stage can substantially influence subsequent analyses, and in the current age of reproducibility crisis, the transparency of this process is vital.
2. Here we present an open-access, reproducible toolkit written in the programming language R for processing raw data files into a single cleaned data set for analyses and upload to online tracking databases (found here: https://github.com/ExMove/ExMove). The toolkit comprises well documented and flexible code to facilitate data processing and user understanding, both of which can increase user confidence and improve the uptake of sharing open and reproducible code. Additionally, we provide an overview website (found here: https://exmove.github.io/) and a Shiny app to help users visualise tracking data and assist with parameter determination during data cleaning.
3. The toolkit is generalisable to different data formats and device types, uses modern ‘tidy coding’ practices, and relies on few well-maintained packages. Amongst these, we perform spatial manipulations using the package *sf*, providing a timely resource for movement ecologists following the recent deprecation of the widely-used *sp* package.
4. Overall, by collating all required steps from data collection to archiving on open access databases into a single, robust pipeline, our toolkit provides a valuable resource for anyone conducting animal movement analyses and represents an important step towards increased standardisation and reproducibility in animal movement ecology.

## 1 Introduction

We live in a golden age of animal-tracking data. Recent technological advancements have reduced the size and cost of telemetry devices and improved their capacities, with more species and individuals tracked over longer periods, allowing researchers to expand the frontiers of ecological research (Bodey et al., 2018; Kays et al., 2015; Nathan et al., 2022; Wilmers et al., 2015). These studies have generated a rapidly expanding volume of data (Gupte et al., 2022; Kays et al., 2022), from an increasing range of device types, with associated formats. This proliferation has complicated the workflow from raw files to a cleaned data set suitable for analysis and sharing, with knock-on impacts for reproducibility. Such pre-analysis steps are universal when working with animal movement data and involve the determination of key filtering parameters to remove data that can influence biological inference from subsequent analysis. Moreover, these steps are time consuming (Dasu & Johnson, 2003), often involving a steep learning curve, and large amounts of time and associated grant funding are lost as unacquainted researchers re-run these steps and, in many cases, attempt to re-invent what should be a standardised and reproducible process.

Numerous software tools have been developed to facilitate animal movement research (Beal et al., 2021; Joo et al., 2020). However, these tools largely focus on evaluating data set suitability (Bjørneraas et al., 2010; Morera-Pujol et al., 2023) and conducting movement analyses of cleaned and/or collated data (Jonsen et al., 2023; Kölzsch et al., 2022). The upstream pre- processing steps required to generate an appropriate cleaned data set have been given scant attention (however see (Gupte et al., 2022). The value of transparent data pre-processing and the accessibility of authentic data sets is important amid the push towards open and reproducible science (Culina et al., 2020; Poisot et al., 2019). Tools to facilitate this move can enhance user’s confidence in sharing code (Gomes et al., 2022), and break down barriers to address the reproducibility crisis facing science (Borregaard & Hart, 2016).

There has been progress in developing tools for pre-processing. Gupte et al., (2022) provides an informative framework outlining the key pre-processing steps for high-throughput animal tracking data, alongside an R package, *atlastools*, to implement these steps. MoveApps (Kölzsch et al., 2022) takes an alternative approach to the problem, providing an online user interface where researchers can upload and explore their own tracking data. This provides a tool to rapidly visualise animal tracking data, especially for researchers who lack programming experience. Both approaches do require additional pre-processing of raw data downloads from tags, for example to combine data from multiple individuals, filter data to deployment periods only, and coerce data into specific formats and column names. Collating well-structured data in this way can be complex (Poisot et al., 2019) and poor confidence in code quality to achieve such tasks has contributed to historically low uptake of journal-mandated code archiving (Baker, 2016). Standardised and reproducible pipelines have been highlighted as a key step towards defensible and robust data science (Borregaard & Hart, 2016; Brunsdon & Comber, 2021), whilst also saving individual researchers time and improving learning (Poisot et al., 2019).

Here, we address this gap by providing a code-based toolkit to process raw tracking files into a clean data set. Establishing a standardised and reproducible pipeline that prepares data for advanced animal movement analyses and archiving in online tracking databases; for example Movebank (Kays et al., 2022) and the Seabird Tracking Database (www.seabirdtracking.org). We have opted for a code-based toolkit shared via versioned GitHub releases rather than an R package, as we want to maximise accessibility and flexibility for a broad user-base. Experienced movement analysts can benefit from rapid data processing, whilst newer users can learn modern R coding skills for processing their tracking data. Our methods involve thorough interrogation of the user-defined parameters for data filtering that can have important downstream consequences for biological inference. By sharing transparent code, we equip users with tools to learn from and adapt for their own analyses. This will make processing tracking data accessible to a wider audience, breaking down financial barriers associated with expensive training courses, while simultaneously flattening the learning curve for ecologists everywhere. The ExMove toolkit aims to equip researchers at all levels with accessible, flexible, and shareable code, and help contribute towards more open-access attitudes in science (Barnes, 2010).

## 2 Objectives

Given the increase in animal movement research, and current gaps in the availability and accessibility of data pre-processing tools, we have developed a toolkit in the programming language R. This manuscript and the code provided have the following overall objectives:

1. Provide a flexible pipeline to process data from tracking devices, such as Global Positioning System (GPS), light-level geolocators (GLS) and Argos tags, into a clean and processed data set upon which researchers can build their own bespoke animal movement analyses.
2. Streamline the path from data collection to sharing of cleaned data sets in online repositories to facilitate open-access and reproducibility requirements of funders, journals, and institutions.
3. Openly-share a resource for users to learn appropriate R coding tools whilst processing animal telemetry data. This will help to develop user confidence in their code quality and promote code-sharing, leading to improved reproducibility within movement ecology (Gomes et al., 2022; Poisot et al., 2019).
4. Facilitate rapid and efficient parameter determination for data processing by providing an interactive Shiny App, which incorporates a range of useful visualizations to explore data.
5. Maximise stability by using a few well-maintained core R packages, including here, tidyverse and sf. Code that implements these packages is especially relevant given the widespread adoption of tidy-style coding and the recent deprecation of *sp* (Müller & Bryan, 2020; Pebesma, 2018; Wickham et al., 2019).

## 3 Workflow

Here, we provide an outline of the code-based workflow from the ExMove toolkit (Figure 1) that enable a user to import raw tracking data and metadata into R and then perform data cleaning, filtering, and processing steps, ready for further analysis. The workflow is flexible and is designed to accommodate anyone undertaking analysis of animal movement data. By following this workflow, users will achieve the following minimum requirements for clean data: date and time formats standardised across time zones, relevant metadata included, no-data values removed (e.g., “NA”, “0,0” in latitude/longitude or other erroneous values), erroneous speeds and locations removed, duplicates removed, locations within pre-/post-deployment window removed, and spatial projections standardised. For a full example of how to use all code within the workflow, please see the user guide (Supplementary Material User guide & https://exmove.github.io). The user guide contains a fully worked example of each step, instructions for accessing resources, and an introduction to the directory structure and data principles that we use to ensure standardisation and reproducibility.

**Figure 1.**
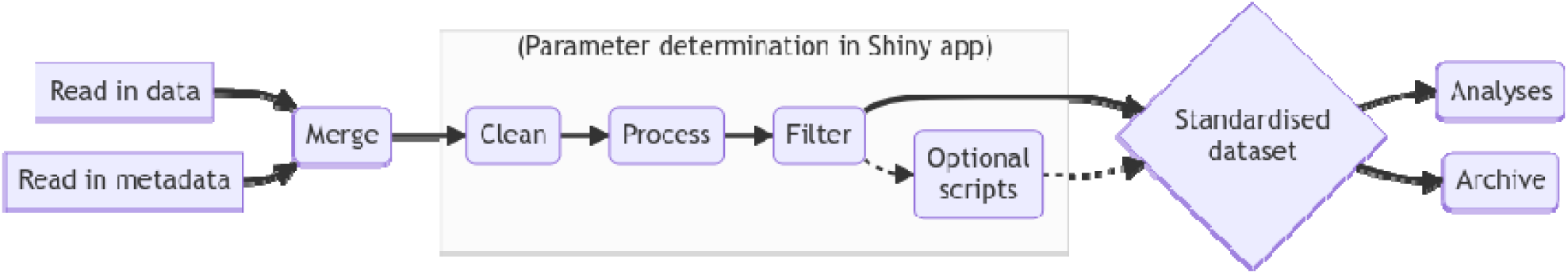
Diagram of workflow used for processing movement data from raw data to archiving (thick line denotes core path of workflow).

### 3.1 Read in data, Read in metadata, and Merge

During the initial step of the workflow, we read in raw tracking data and metadata files and merge them (user guide sections 1-2). Raw tracking data refers to data downloaded directly from telemetry devices, which fundamentally must contain a time stamp and location data. We also provide alternative code for data that have already been partially processed, for example data downloaded from an existing repository that contains multiple tags and/or individuals. Metadata provide essential technical, biological, and ecological context for the raw data, such as the tag deployment period, sex, age and population. Metadata should contain one row for each tag deployment per individual, with ID numbers corresponding to those in the raw data. The ID numbers are used to match each row of the raw data to the corresponding metadata. Additional code is provided to handle scenarios where a single individual has carried multiple tags in its lifetime, or a single tag has been deployed on multiple individuals. Example data are saved as .csv files (see supplementary files), and guidance is provided for importing data in other text-based formats (e.g., .txt or .tsv). In addition to telemetry data, the code from this step can also be adapted for biologging studies, to merge and standardise additional sensor data (e.g., water immersion or depth) from multiple individuals.

### 3.2 Cleaning

In the cleaning step, we remove data that are incorrect based on deployment periods and technical tag information (user guide section 3). This includes limiting data to the deployment period (i.e., for loggers programmed to start ahead of deployment or not turned off immediately following retrieval), removing duplicate rows, and removing NA’s or known values that correspond to missing data based on tag manufacturer guidance. Some tags record a specific ‘no data’ value when sensor information cannot be derived (e.g., “0,0” for latitude and longitude).

### 3.3 Processing

During the processing step we calculate spatial and temporal metrics to allow downstream analyses and filtering of data based on ecological knowledge and device configuration (user guide section 4). These include metrics pertaining to ecological processes, such as distance travelled (Euclidian distance between successive locations), net displacement (Euclidian distance from first location for each unique ID, e.g., used to quantify migration phenology; (Soriano-Redondo et al., 2020), travel speed (distance travelled dived by time step duration), and turning angle (trajectory of movement at current time step relative to previous trajectory). We also include time step duration (duration between successive locations), which can vary depending on device configuration and performance, but is informative for downstream movement analyses. See Fig. 3 for graphical visualization of these metrics, the code for which is presented in the workflow and incorporated into the shiny app.

**Figure 2.**
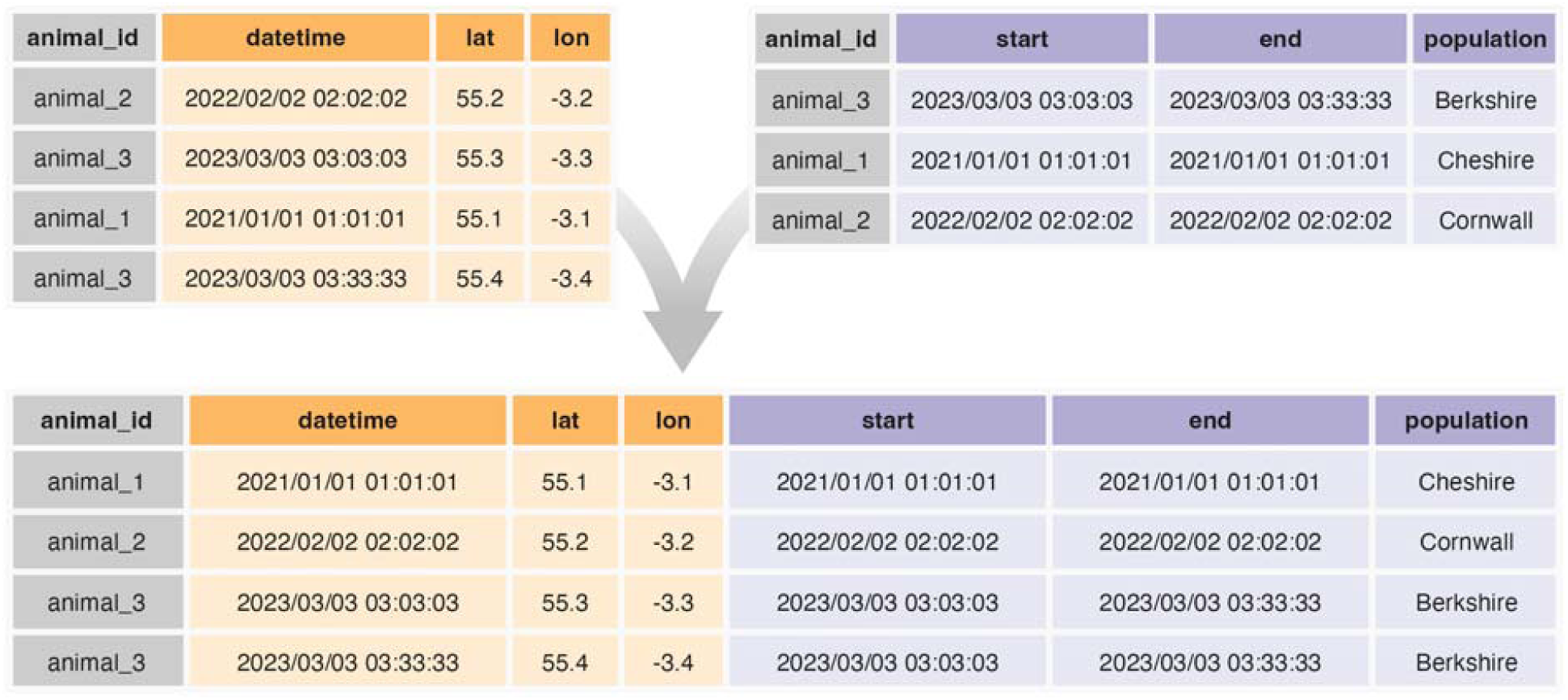
Example of combining raw GPS tracking data file (orange) and metadata file (lilac), using the ExMove toolkit, with “animal_id” as the index column. Note that accuracy of latitude and longitude data has been reduced for clarity.

**Figure 3.**
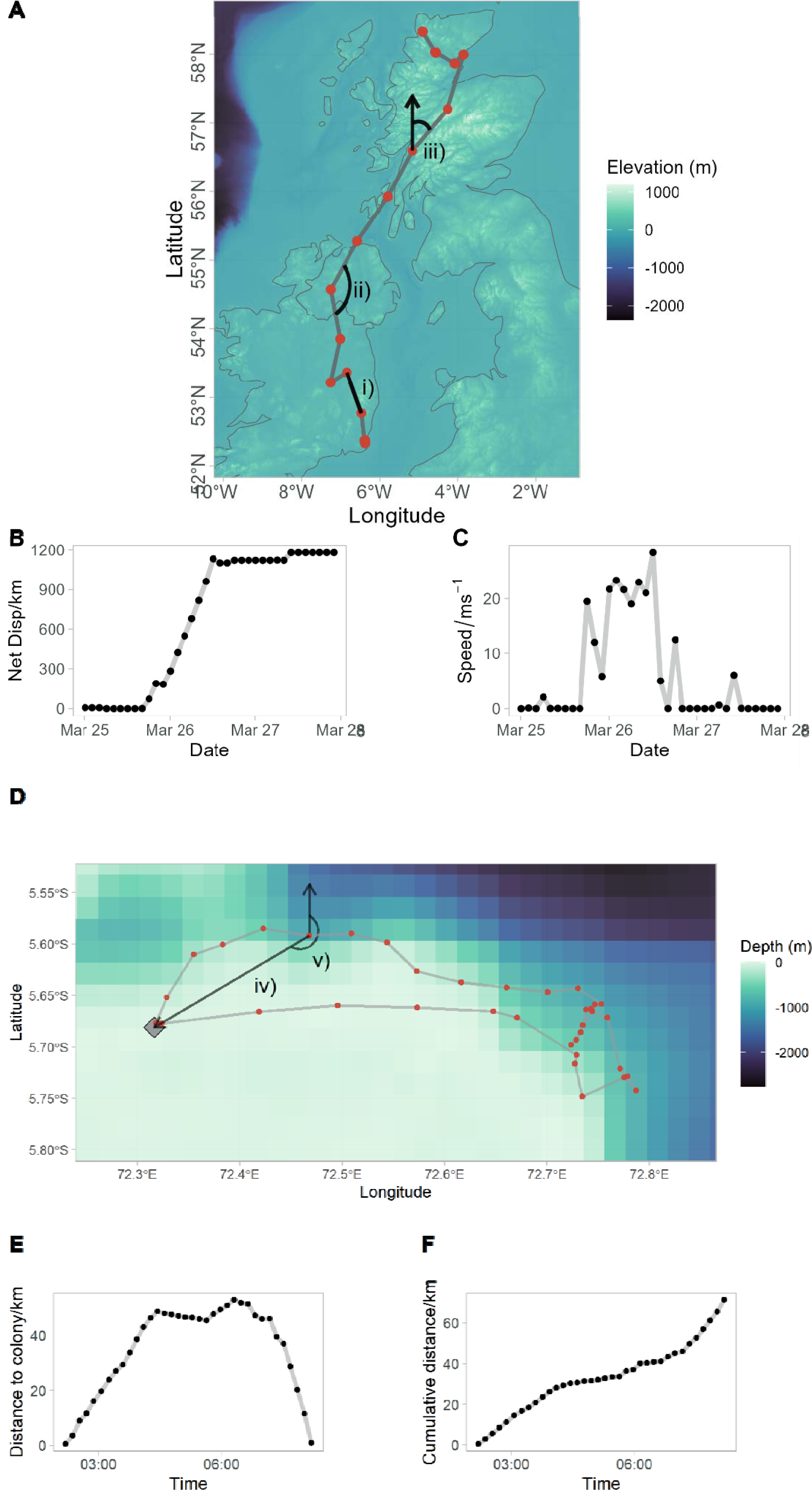
Processed metrics derived from animal tracking data. Plot A depicts a movement track of a Greenland White-fronted Goose with three different metrics labelled: i) step length, ii) turning angle and iii) travel bearing. The net displacement (plot B) and speed (plot C) over time are shown for the movement track in plot A. Plot D depicts a single foraging trip of a Red-footed Booby from a central place (grey filled diamond), illustrating the metrics: iv) distance to central place and v) bearing to central place for a given location. The distance to colony (plot E) and cumulative distance travelled (plot F) are shown for the track in Plot D.

Location information from biologging devices is recorded in a specific Coordinate Reference System (CRS). Most often, locations are recorded as degrees of latitude and longitude (CRS: WGS84) which are ‘unprojected’ representations of locations on a sphere, and are generally not recommended for performing distance calculations. ‘Projected’ spatial data (like the format we convert data to in the workflow) are characterised by a CRS that refers to the 2D projection of the earth’s surface used to represent locations. This type of CRS is recommended for calculating distances and angles and is defined by the user during the processing step. We transform data in the workflow to a global ‘equal area projection’ for spatial calculation, which can be useful for tracking data spanning a large geographic range. However, there is no “correct” CRS and the specific choice may depend on whether users want to preserve distances or angles, and the spatial extent of the data. We provide further guidance and links in section 4 of our FAQs (https://exmove.github.io/FAQs.html).

### 3.4 Filtering

During the filtering step we remove erroneous locations based on three derived variables: 1) speed, 2) net displacement and 3) time since tag deployment (user guide section 6). Each of these variables are described and calculated during the 3.3 processing step (Fig 3). Threshold values are selected by the user to filter the data – if these values are exceeded then the containing rows are removed. The downstream consequences of filtering decision can be explored with the visualizations we provided in our Shiny app (see 4.3). Ecological knowledge from the user is required to help inform thresholds: for instance a species’ maximum travel speed, or the time after tag deployment that behaviour is expected to return to normal. We encourage users to explore a range of values and visualise the consequences using maps of movement tracks and histograms of speed and net squared displacement.

### 3.5 Summarise

Here we provide a summary of the data that the user has retained after the filtering step (user guide section 7). This is an important opportunity to scrutinise the integrity of the data and determine if further filtering is required. The user selects grouping variables for the data — we suggest initially grouping by individual, but broader (e.g., year and population) or narrower (e.g. date) groupings may be required depending on the type of analysis to be undertaken. For each level of the grouping variable, the following summary statistics are provided: number of locations, number of unique tracking days, start and end dates of tracking period, and the sampling frequency locations.

### 3.6 Post-processing: segmenting, sub-sampling & classifying foraging trips (optional)

Some types of animal movement analyses require additional processing to prepare data (e.g., hidden Markov models (Langrock et al., 2012) and step selection functions (Fieberg et al., 2021)). The toolkit includes code to implement four types of optional processing step: segmenting, sub-sampling, classifying foraging trips and track animation. Segmenting separates tracks into bursts of continuous data with consistent sampling intervals to avoid over-interpolation across large temporal gaps. Tracks are segmented into separate bursts if the time between two locations exceeds a user defined threshold. Then, only bursts that contain a minimum number of locations are kept for further analysis and this threshold is also defined by the user. Sub-sampling aims to standardise the sampling interval across a track by thinning the data, thus facilitating comparisons among data sets with different original sampling regimes (Fig 4). The user defines a new temporal sampling frequency which should be lower than the average sampling frequency determined at earlier steps (i.e., greater time steps). In cases where increasing sampling frequency, or interpolation, are required, we recommend users to find appropriate methods for their study system and question. For example, data exported from the ExMove toolkit is in an appropriate format for the *aniMotum* package, which uses error radii around telemetry locations to derive most likely movement trajectories (Jonsen et al., 2023).

**Figure 4.**
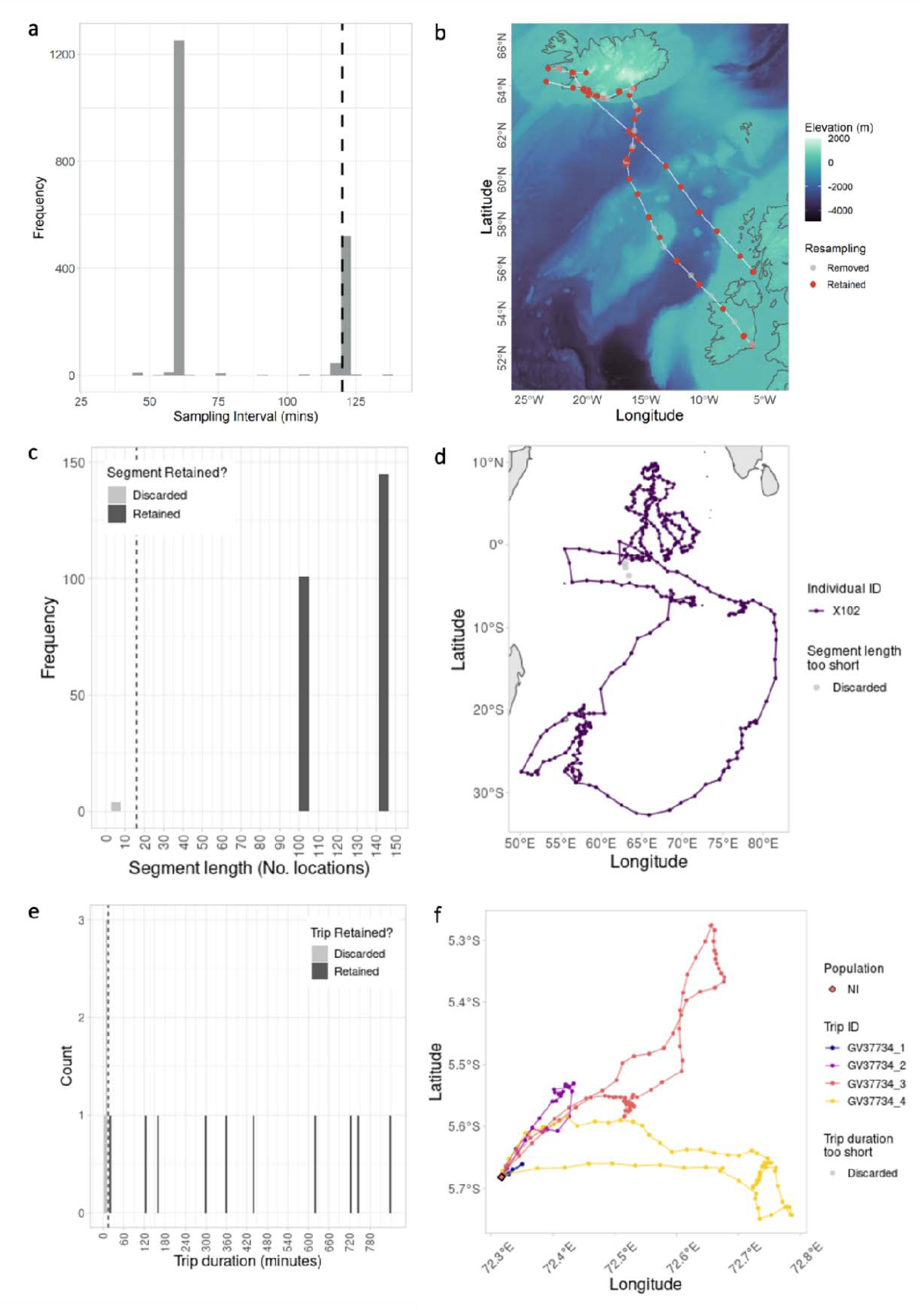
Example of optional post-processing scripts to (a-b) sub-sample data to lower temporal resolution, (c-d) identify segments split by longer temporal intervals, and (e-f) define central-place foraging trips. (a-b) sampling intervals in Greenland White-fronted Geese data include data collected at one- and two-hour resolutions have been sub-sampled to 2-hour resolution (dashed line in panel a) to standardise data across individuals (red: retained, grey: removed fixes in panel b). (c-d) short segments, containing fewer than 18 tracking locations (dashed line in panel c) have been discarded from the migration track of a Trindade petrel (purple: retained, grey: removed fixes in panel b). (e-f) short trips, less than 15 minutes duration (dashed line in panel e) have been discarded from the central-place foraging trips of a red-footed booby (grey points in panel f are less than 1km from the colony, and so have been obscured by retained trips in distinct colours).

For central place foragers, we include an optional script to calculate distance and bearing to a central place (typically a breeding colony or roosting site) and classify distinct foraging trips (Trevail et al., 2019). The central place is user-defined and may refer to a known individual nest site, a central colony location for the population, or a shapefile/polygon delineating the spatial boundary of the central place. We provide code to classify and number trips away from the central place using either a distance threshold approach or based on birds moving outside the colony area boundary, and then to classify trips as ‘foraging’ trips using a time threshold, facilitating the removal of very short trips (e.g., in response to disturbance events). We then calculate standard metrics of foraging trips: total distance travelled, maximum distance from the central place, trip duration, and distal location, to facilitate standardised reporting of central place forager’s behavioural ecology. Finally, we include a script to animate tracks allowing users to dynamically visualise movement trajectories of their organisms over time.

All post-processing steps are provided as additional scripts and are designed to follow on from the main code workflow. Scripts can be run iteratively if multiple additional post-processing steps are required. Each script contains tools to summarise, visualise, and save the post-processed data. The Shiny app can also be used here to aid parameter determination.

### 3.7 Standardising output for tracking databases

Open and reproducible science requires data to be presented and archived in a standard, replicable, interoperable way. Online tracking data repositories have achieved this by following the FAIR principles (Wilkinson et al., 2016) and the TRUST principles (Lin et al., 2020). We encourage archiving of tracking data in shared online repositories and as such, we provide code for users to format and save their data, processed within the workflow, in a standardised format that is accepted by two common tracking databases: (1) MoveBank (Kays et al., 2022), and (2) the Seabird Tracking Database.

### 3.8 Toolkit Continuation

We hope to see widespread adoption of the ExMove toolkit as a starting point for standardised cleaning and processing of animal telemetry data. If users encounter bugs within the toolkit or have suggestions regarding enhancements we encourage them to get in contact with us by raising issues or pull requests via the GitHub. If we make future large-scale changes or add extra functionality we will issue an update via a future GitHub release.

## 4 Tools and additional practical resources

### 4.1 User guide

We provide a fully-coded user guide that follows each step of the workflow with worked examples in tidy code. This guide contains example data outputs, plots produced for each step, and detailed annotations to help users adapt the code to their data set. The user guide is available in supplementary material, within the GitHub repository (https://github.com/ExMove/ExMove/tree/main/Documentation), and via the ExMove website (https://exmove.github.io/User_guide.html).

### 4.2 FAQ’s

Some important principles of movement and spatial analyses can be a common source of confusion (for example: understanding data projections). Given ExMove’s goal as a learning tool, we provide information to help facilitate better understanding of these core concepts, and links to useful materials made available by others (FAQ page can be found here: https://github.com/ExMove/ExMove/tree/main/Documentation & https://exmove.github.io/FAQs.html).

### 4.3 Shiny app

The Shiny app is available online at the following link: https://lukeozsanlav.shinyapps.io/exmove_explorer/. The shiny app follows the same steps as the workflow in sections 3.4 to 3.6 (section 6 onwards of user guide) but they are incorporated into a user friendly user interface. The code for the shiny app is also available in the GitHub repository, alongside the workflow. There is an upload limit so users can either explore the app with examples data sets we provide or a subset of their own data. If the user wants to run the app on their full data set, then we recommend that they clone the repository and then run the app locally on their own machine. The code for doing this is explained in section 6 of the user guide.

The app is designed to be accessible for new R users to visualise and filter data but also provides a convenient interactive platform for more experienced users to determine the downstream effects of user-input parameters on the resulting cleaned data frames. When processing animal tracking data, prior to analysis, users often select rather arbitrary values by which to filter data, i.e., to eliminate erroneous speeds users often remove points creating speeds above a selected threshold. The app allows users to explore the implications of these arbitrary decisions via interactive manipulation of threshold values and subsequent visualizations that include, data summary tables, histograms of the variable of interest and maps. There are multiple tabs to navigate within the app once users have uploaded their own data, in each there are a series of interactive user input threshold values and a number of plots and tables detailing the consequences of the user selected values. There are tabs for basic data filters (e.g., speed, post-deployment period and date), calculating foraging trips for central placed foragers, sub-sampling location frequency, segmenting movement tracks and finally visualizing the locations in an interactive leaflet map. To facilitate integration with the main workflow there are code chunks output at the bottom of each shiny tab that contain the selected parameter values, which can be copied into the workflow.

### 4.4 Example data sets

Within the GitHub repo we provide four tested example data sets for users to familiarise themselves with using the toolkit. These data sets are from Trindade Petrels (*Pterodroma arminjoniana*; Figure 5A) breeding on Round Island (Nicoll et al., 2017), immature and adult red-footed boobies (*Sula leucogaster*, Figure 5B & D) breeding in the Chagos Archipelago (Trevail et al 2023) and Greenland white-fronted goose (*Anser albifrons flavirostris*, Figure 5C) migrating from Ireland and Scotland to Greenland (Ozsanlav-Harris et al., 2022). These data sets were selected to illustrate how our toolkit is able to process telemetry data from different tag types, species, life-history stages and spatial movement scales. They also illustrate how running telemetry data through our optional processing scripts can provide additional ecological insight, such as defining foraging trips for central-place foragers (Fig 5D & Fig 6). An example processing pipeline for central-place foraging red-footed boobies is shown in Figure 6, for which all code is fully presented and described in the user guide.

**Figure 5.**
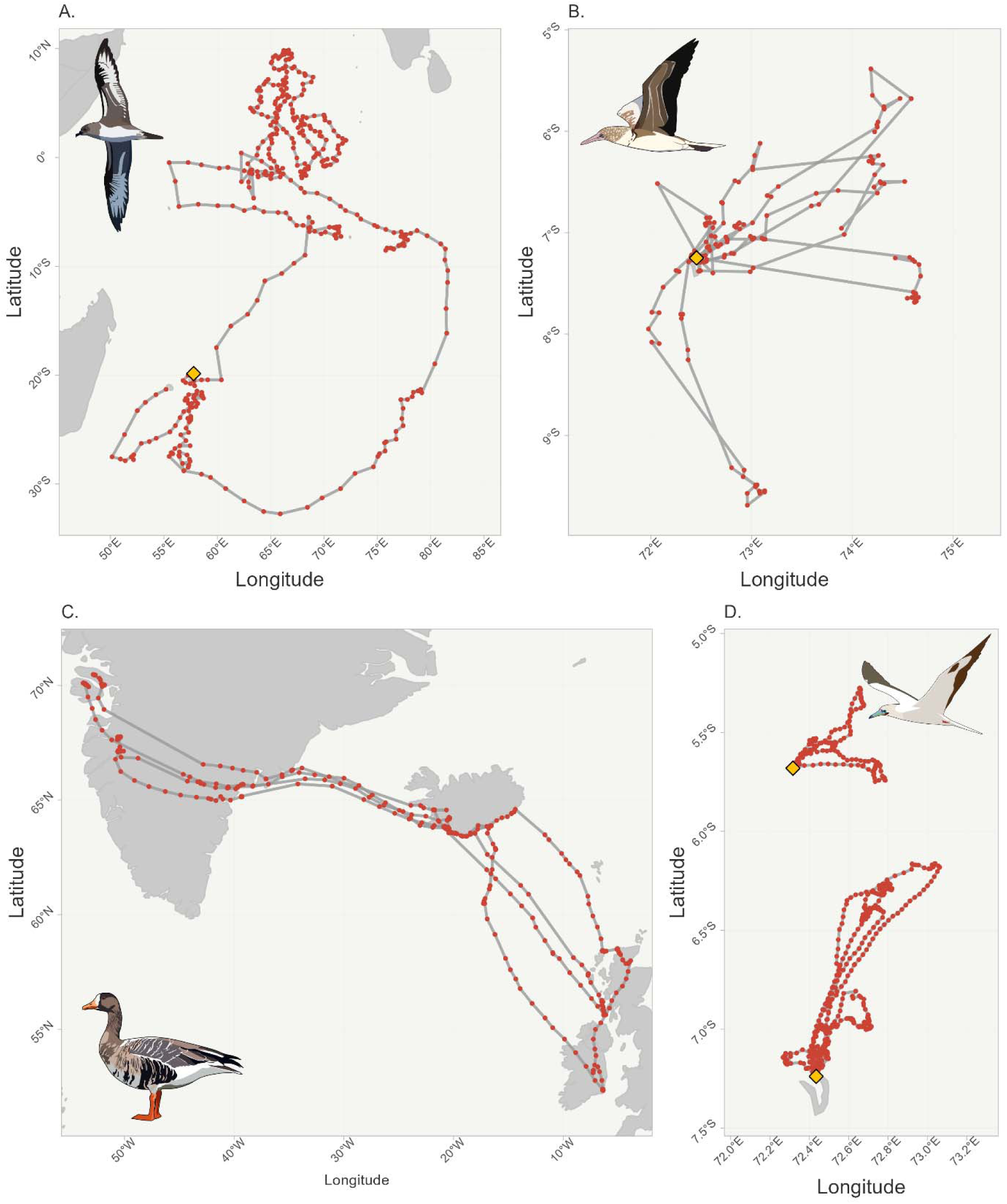
Visualisation of the four example data sets that have been cleaned by the toolkit, with individual telemetry points (red dots) and breeding colony locations (orange diamonds). A: GLS data from a Trindade petrel during its non-breeding migration in 2011. The device was deployed and retrieved from Round Island Nature Reserve (19.85° south; 57.78° east), situated 23 km off the North coast of Mauritius, Indian Ocean. For methods and ethics see Nicoll et al., (2017). B: Argos data from two immature red-footed boobies tracked during 2022 using Microwave Telemetry Solar Argos/GPS PTT loggers (Figure 6B). Data represent foraging movements of immature individuals tagged whilst roosting on East Island in the Chagos Archipelago, Indian Ocean. C: GPS data from four Greenland White-fronted Geese tracked during spring migration in 2018 and 2019. For methods and ethics see Ozsanlav-Harris *et al*. (2022). D: GPS data from three adult red-footed boobies tracked during 2016 and 2018 using igotU loggers (Figure 6D). Data represent central-place foraging trips of breeding adults from two colonies, Diego Garcia and Nelsons Island in the Chagos Archipelago, Indian Ocean. For methods and ethics see Trevail *et al*. (2023).

**Figure 6.**
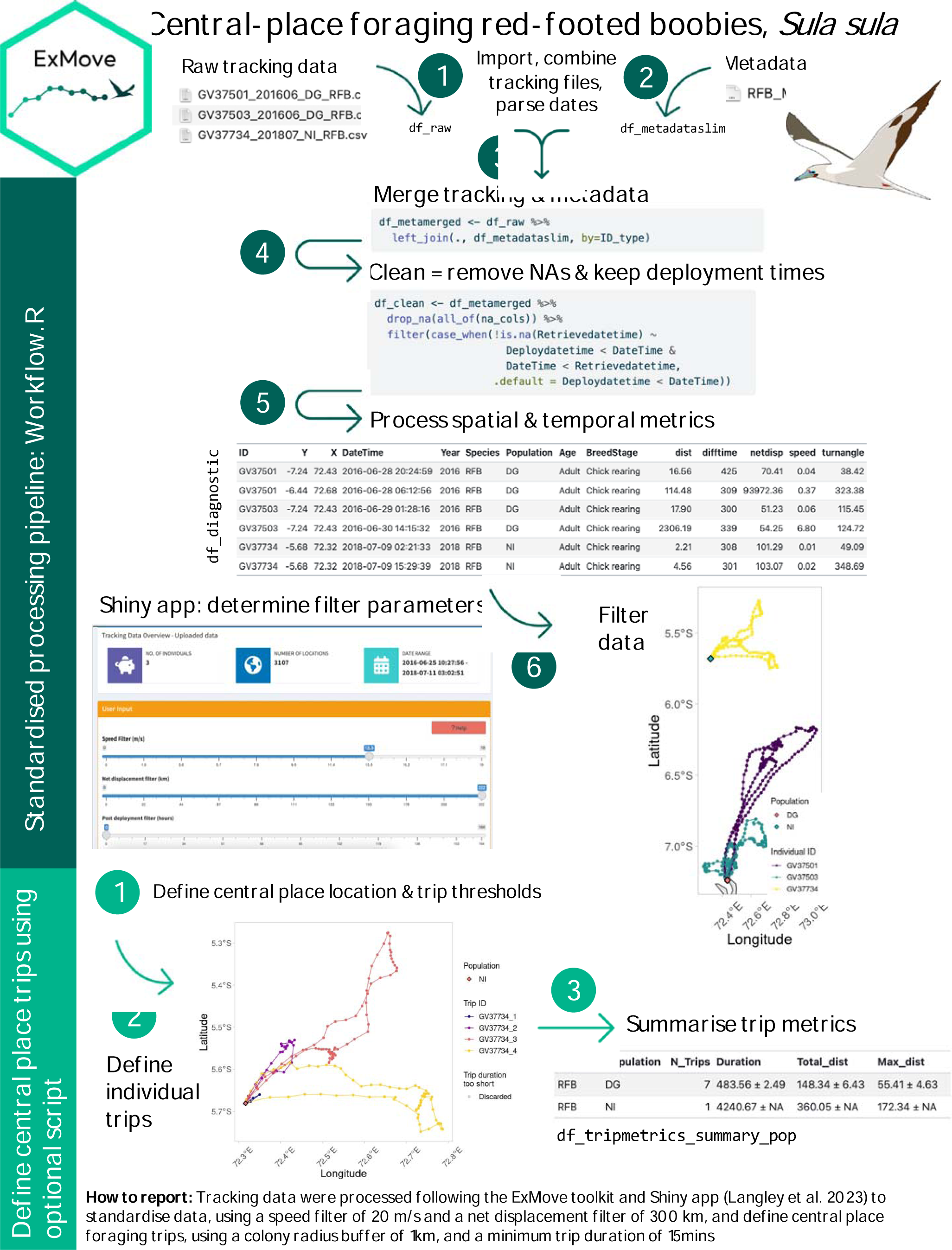
An example of how the ExMove toolkit has been applied to GPS-tracked central-place foraging red-footed boobies, *Sula sula.* The toolkit facilitates import of raw data and metadata, cleaning, processing, and filtering within the ‘Workflow.R‘ script. Then identification and summary of central-place behaviour in the ‘Optional Processing_Central place trips.R‘ script. The Shiny app aids parameter determination for filtering (speed and net-displacement) and central-place trip definition (distance from colony and minimum trip duration). Results from the tracking study are presented within Trevail et al. (2023).

## Supporting information

Supplementary Material User guide

## Acknowledgments

Alice Trevail was funded by the Bertarelli Programme in Marine Science, who also funded collection of adult and immature red-footed booby data. Stephen Lang was funded by the Pew Charitable Trusts. We thank the Wildfowl and Wetlands Trust for providing tracking data from Greenland White-fronted Geese and Larry Griffin for deploying tags. We thank Malcolm Nicoll for valuable comments on an earlier version of the manuscript, and for providing tracking data from a Trindade petrel. The Trindade petrel tracking programme (Round Island, Mauritius) was supported by NERC (Grant NE/H5081500) and Research England, with thanks to Mauritian Wildlife Foundation (MWF) and the National Parks and Conservation Service (NPCS). We also thank Evelyn Alexander, Jacqueline Glencross and Tess Handby for providing valuable feedback on the code, workflow and Shiny app.

## Author contributions

All authors contributed equally to the manuscript.

## Data availability statement

All data and code presented in this manuscript can be found in the following public GitHub repository release “ExMove 1.0.0” created at the time of publication (GitHub release TBC): https://github.com/ExMove/ExMove. All tracking data sets used are cited appropriately in section 4.4.

## Notes

https://github.com/ExMove/ExMove

### Competing Interest Statement

The authors have declared no competing interest.

### Summary of Updates

Second Revisions. The overall flow of the manuscript has been improved as well as the accompanying user guide.

