## Supplementary Material User guide for "ExMove: An open-source toolkit for processing and exploring animal tracking data in R"

### ExMove user guide

Code

#### Introduction

This user guide can be used as a walkthrough for reading and processing tracking data files with the `Workflow.R` script. You can use the example datasets provided in `Data`, or try with your own tracking data (see Pre-flight checks for details on data requirements and structure).

The following diagram gives an overview of the workflow (boxes link to relevant section):

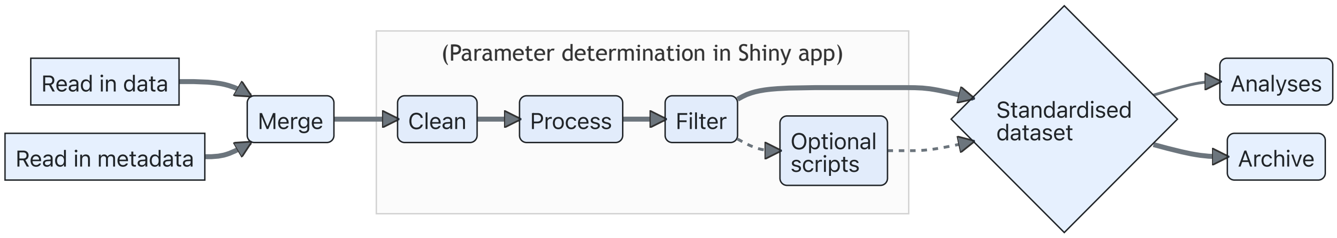


Figure 1: Diagram of workflow used for analysing movement data (thick line denotes core path of code)

##### Dependencies:

- This workflow uses the R programming language, run via the R Studio IDE
- All code embraces the core principles of how to structure 'tidy data'
- We use RStudio projects and the `here` package to build relative filepaths that are reproducible
- Requires `tidyverse`, `data.table`, `sf` and `here` packages to be installed
- Use our example data sets in the Data folder (RFB_IMM, RFB, GWFG, TRPE) or provide your own data

##### User inputs

Some code chunks require editing by the user to match the specific dataset being used (particularly if you are using your own data), and are highlighted as below (the 🧠 indicates you will need to think about the structure and format of your data when making these edits!):

🧠 **User input required**

```
#--------------------#
## USER INPUT START ##
#--------------------#
example_input <- "uservalue" # In the R code, user input sections appear like this
#------------------#
```

```
## USER INPUT END ##
#-----------------#
```

### 0. Pre-flight checks

##### How to use this workflow:

- We will inspect the data before reading it in, so there is no need to open it in another program (e.g., excel, which can corrupt dates and times)
- User-defined parameters (see user inputs) are called within the subsequent processing steps
- Where you see: `## ** Option ** ##`, there is an alternative version of the code to fit some common alternative data formats
- Throughout, we will use some key functions to inspect the data (e.g., `head` for top rows, `str` for column types, and `names` for column names)

##### Data and directory structure:

- Data files should all be stored within the `Data` folder
- Folders and files are best named in `snakecase_format` as spaces in filepaths can cause issues
- Tracking data for each deployment/individual should be in a separate file
- Tracking data filenames should include an ID which is the same length for all individuals
- Tracking data must contain a timestamp and at least one other sensor column
- Metadata file should be in the parent directory of data files
- Metadata should contain one row per individual per deployment *NB: if you have multiple animal movement projects, these should have completely separate directories*

##### The importance of ID:

- Throughout this workflow, we use ID to refer to the unique code for an individual animal
- In certain cases, you might have additional ID columns in the metadata (e.g., `DeployID`),
- or read in data with a unique TagID instead of ID.
- This code will work as long as all of the relevant info is included in the metadata
- For more info and helpful code, see the FAQ document & troubleshooting script

##### How to troubleshoot problems if something doesn't work with your data:

- Refer to the FAQ document in the GitHub page, which signposts to helpful resources online (e.g., CRS)
- See the troubleshooting code scripts that we've written to accompany this workflow (e.g., using multiple ID columns for re-deployments of tags/individuals)
- All functions in code chunks are automatically hyperlinked to their documentation, so feel free to explore this if you want to understand more about how this code works!

##### Load required libraries

Just before starting we load in all the packages we will need for the workflow (also referenced in the Dependencies section).

```
library(data.table) # data manipulation
library(tidyverse) # data reading, manipulation and plotting
library(lubridate) # working with date-time data
```

```
library(sf) # spatial data handling and manipulation
library(here) # reproducible filepaths
```

### 1. Read in data files

🧠 **User input required**

Throughout the script, we'll be saving files using a species code as a file/folder identifier. Our code is taken from the first letters of the species name (**R**ed **F**ooted **B**ooby) but can be anything you choose (shorter the better as long as it's unique per species). Let's define this object first for consistency:

```
species_code <- "RFB"
```

Set filepath for the folder containing raw data files (this code will try to list and open all files matching the file pattern within this folder, so it is best if this folder contains only the raw data files). ***NB:*** *if you are working outside of a project, you'll need to check that here is using the correct working directory.*

```
filepath <- here("Data", species_code)  #create relative filepath using folder name and specie
```

Define common file pattern to look for. An asterisk (`*`) is the wildcard, will match any character except a forward-slash (e.g. `*.csv` will import all files that end with ".csv").

```
filepattern <- "*.csv" # data file format
```

Let's view the file names, to check that we have the files we want & find ID position (this list will include names of sub-folders).

```
ls_filenames <- list.files(path = filepath, recursive = TRUE, pattern = filepattern)
ls_filenames
```

```
[1] "GV37501_201606_DG_RFB.csv" "GV37503_201606_DG_RFB.csv"
[3] "GV37734_201807_NI_RFB.csv"
```

Adjust these numbers for extracting the ID number from file name using `stringr` (e.g. to extract `GV37501` from "GV37501_201606_DG_RFB.csv", we want characters 1-7). ***NB:*** *this approach only works if all ID's are the same length and in the same position — see the `str_sub` documentation for other options.*

```
IDstart <- 1 #start position of the ID in the filename
IDend <- 7 #end position of the ID in the filename
```

Now, let's inspect the data by reading in the top of the first data file as raw text. To inspect the first row of all data files (if you wanted to check column names), you can remove the `[1]` and change `n_max = 1`).

```
test <- fs::dir_ls(path = filepath, recurse = TRUE, type = "file",  glob = filepattern)[1]
read_lines(test, n_max = 5)  # change n_max to change the number of rows to read in
```

```
[1] "Date, Time, Latitude, Longitude, Altitude, Speed, Course, Type, Distance"
[2] "24/06/2016,5.06,-7.261829,72.376091,56.18,3636,93,-2,0.00"
[3] "24/06/2016, 05:21:39,-7.261829,72.376091,56.18,3636,93,0,0.00"
[4] "24/06/2016, 05:29:50,-7.261744,72.376221,64.91,5112,32,0,17.17"
[5] "24/06/2016, 05:54:28,-7.261623,72.376137,-15.77,936,291,0,16.34"
```

Define number of lines at top of file to skip (e.g. if importing a text file with additional info at top).

```
skiplines <- 0
```

Define date format(s) used (for passing to lubridate) (`d` = day as decimal, `m` = month as decimal, `y` = year without century, `Y` = year with century). Parsing will work the same for different date delimiters (e.g. "dmY" will work for both `01-12-2022` and `01/12/2022`). `lubridate` can even parse more than one date/time format within a dataframe, so if your data include multiple formats, make sure they are all included. Here, we've included some common combinations — modify if your data include a different format

```
date_formats <- c("dmY", "Ymd") #specify date formats
datetime_formats <- c("dmY HMS", "Ymd HMS") #specify date & time format
```

Define time zone for tracking data.

```
trackingdatatimezone <- "GMT"
```

By default, the below code will find column names from the first row of data. If you want to specify your own column names, do so here as a character vector, or use `set colnames <- FALSE` to automatically number columns.

```
colnames <- TRUE
```

Here, we use the function `read_delim` and specify the delimiter to make this code more universal (you can find extra information on this in the readr documentation).

Some delimiter examples:

- `","` = comma delimited (equivalent to using `read_csv` – saved as extension `.csv`)
- `"\t"` = tab delimited (equivalent to using `read_tsv` — saved as extension `.tsv`)
- `" "` = whitespace delimited (equivalent to using `read_table`)

Let's inspect the data again, this time skipping rows if set, to check the file delimiter.

```
read_lines(test, n_max = 5, skip = skiplines)
```

```
[1] "Date, Time, Latitude, Longitude, Altitude, Speed, Course, Type, Distance"
[2] "24/06/2016,5.06,-7.261829,72.376091,56.18,3636,93,-2,0.00"
[3] "24/06/2016, 05:21:39,-7.261829,72.376091,56.18,3636,93,0,0.00"
[4] "24/06/2016, 05:29:50,-7.261744,72.376221,64.91,5112,32,0,17.17"
[5] "24/06/2016, 05:54:28,-7.261623,72.376137,-15.77,936,291,0,16.34"
```

Set delimiter to use within `read_delim`.

```
user_delim <- ","
user_trim_ws <- TRUE # Should leading/trailing whitespaces be trimmed
```

Finally, data need an ID column, either be the tag ID ("TagID") or individual ID ("ID"). Specify ID type here, for later matching with the same column in the metadata:

```
ID_type <- "ID"
```

#### Read in and merge all tracking data files

Merge using ID in filename     Option: Merge using ID already in column

With the user inputs specified in the previous section, we'll now read in and merge all tracking data files directly from the github repository, extracting the ID from the filename of each file.

```
df_combined <- fs::dir_ls(path = filepath, # use our defined filepath
                          glob = filepattern, # use file pattern
                          type = "file",  # only list files
                          recurse = TRUE # look inside sub-folders
                          ) %>%
  purrr::set_names(nm = basename(.)) %>% # remove path prefix
  purrr::map_dfr(read_delim, # use read_delim function
                 .id = "filename", # use filename as ID column
                 col_types = cols(.default = "c"), # as character by default
                 col_names = colnames, # use colnames object made above
                 skip = skiplines, # how many lines to skip
                 delim = user_delim, # define delimiter
                 trim_ws = user_trim_ws) %>% # trim characters or not
  mutate("{ID_type}" := str_sub(string = filename, # extract ID from filename
                                start = IDstart, end = IDend), # ID position
         .after = filename) # move the new ID column after filename column
df_combined
```

```
# A tibble: 6,650 × 12
   filename     ID    Date  Time  Latitude Longitude Altitude Speed Course Type
   <chr>        <chr> <chr> <chr> <chr>    <chr>     <chr>    <chr> <chr>  <chr>
 1 GV37501_201… GV37… 24/0… 5.06  -7.2618… 72.376091 56.18    3636  93     -2
 2 GV37501_201… GV37… 24/0… 05:2… -7.2618… 72.376091 56.18    3636  93     0
 3 GV37501_201… GV37… 24/0… 05:2… -7.2617… 72.376221 64.91    5112  32     0
 4 GV37501_201… GV37… 24/0… 05:5… -7.2616… 72.376137 -15.77   936   291    0
 5 GV37501_201… GV37… 24/0… 06:2… -7.2616… 72.376335 30.91    1080  124    0
 6 GV37501_201… GV37… 24/0… 06:2… -7.2616… 72.376289 26.8     0     280    0
 7 GV37501_201… GV37… 24/0… 06:3… -7.2615… 72.376198 21.77    0     183    0
 8 GV37501_201… GV37… 24/0… 06:3… -7.2615… 72.376205 30.78    2844  268    0
 9 GV37501_201… GV37… 24/0… 06:4… -7.2616… 72.376259 27.1     1476  44     0
10 GV37501_201… GV37… 24/0… 06:4… -7.2614… 72.375732 -35.04   0     260    0
# ℹ 6,640 more rows
# ℹ 2 more variables: Distance <chr>, Essential <chr>
```

```
colnames(df_combined)
```

```
 [1] "filename"  "ID"        "Date"      "Time"      "Latitude"  "Longitude"
 [7] "Altitude"  "Speed"     "Course"    "Type"      "Distance"  "Essential"
```

#### Slim down dataset

Select normal columns     Option: Select custom columns

🧠 User input required

First, data need a time stamp, either in separate columns (e.g., “Date” and “Time”) or combined (“DateTime”). Below we specify which column’s date and time info are stored in the data. ***NB:*** *These have to be in the same order as specified in earlier user input, i.e. “Date” and “Time” have to be the right way round*

```
datetime_formats # previously specified datetime formats
```

```
[1] "dmY HMS" "Ymd HMS"
```

```
datetime_colnames <- c("Date", "Time") # or c("DateTime")
```

You can also have additional columns depending on the type of logger used, for example:

```
## lc = Argos fix quality
## Lat2/Lon2 = additional location fixes from Argos tag
## laterr/lonerr = location error information provided by some GLS processing packages
```

Here we’re going to slim down the dataset by selecting the necessary columns & coercing some column names. You should change column names below to those present in your tracking data, additional columns can be added (see above examples). This process standardises important column names for the rest of the workflow (e.g., `TagID`, `Lat`, `Lon`)

```
df_slim <- data.frame(ID = as.character(df_combined$ID),
                      Date = df_combined$Date,
                      Time = df_combined$Time,
                      Y = df_combined$Latitude,
                      X = df_combined$Longitude)
```

#### Parse dates, create datetime, date and year columns

Now our `df_slim` is ready, we need to create a `DateTime` column. Using the `datetime_colnames` object we made previously, we’ll combine columns (if needed), and then parse a single `DateTime` column using the `lubridate` package:

```
df_slim <- df_slim %>%
  tidyr::unite(col = "DateTime_unparsed", # united column name
               all_of(datetime_colnames), # which columns to unite
               sep = " ",  # separator between values in new column
               remove = FALSE # remove original columns?
               ) %>%
  mutate(DateTime = lubridate::parse_date_time(DateTime_unparsed, # parse DateTime
                                               orders = datetime_formats, # formats
                                               tz = trackingdatatimezone), # timezone
         Date = lubridate::as_date(DateTime),
         Year = lubridate::year(DateTime)) %>%
  select(-DateTime_unparsed)
```

```
Warning: There was 1 warning in `mutate()`.
ℹ In argument: `DateTime = lubridate::parse_date_time(...)`.
Caused by warning:
!  1 failed to parse.
```

> **Note**
>
> `n failed to parse` warnings means a date or time was not in the correct format for `lubridate` to create a date_time object, producing `NA`s. We can look at the failing rows using the following code:
>
> ```
> Fails <- df_slim %>% filter(is.na(DateTime)==T)
> head(Fails)
> ```
>
> ```
>        ID Date Time         Y         X DateTime Year
> 1 GV37501 <NA> 5.06 -7.261829 72.376091     <NA>   NA
> ```
>
> Now we can see the issue: `Date` is empty, and `Time` is saved as a number. We'll remove this row in the @cleaning section, so don't need to do anything else for the moment.

Lastly, we make a `df_raw` dataframe by sorting using ID and DateTime, dropping NA's in DateTime column

```
df_raw <- df_slim %>%
  arrange(across(all_of(c(ID_type, "DateTime")))) %>%
  drop_na(DateTime) #remove NA's in datetime column
head(df_raw)
```

```
       ID       Date     Time         Y         X            DateTime Year
1 GV37501 2016-06-24 05:21:39 -7.261829 72.376091 2016-06-24 05:21:39 2016
2 GV37501 2016-06-24 05:29:50 -7.261744 72.376221 2016-06-24 05:29:50 2016
3 GV37501 2016-06-24 05:54:28 -7.261623 72.376137 2016-06-24 05:54:28 2016
4 GV37501 2016-06-24 06:22:02 -7.261651 72.376335 2016-06-24 06:22:02 2016
5 GV37501 2016-06-24 06:27:16 -7.261618 72.376289 2016-06-24 06:27:16 2016
6 GV37501 2016-06-24 06:32:27 -7.261584 72.376198 2016-06-24 06:32:27 2016
```

We can clean up intermediate files/objects by listing everything we want to keep (i.e. remove everything else)

```
rm(list=ls()[!ls() %in% c("df_raw",
                          "date_formats","datetime_formats","trackingdatatimezone",
                          "ID_type", "species_code")])
```

#### 2. Merge with metadata

Metadata are an essential piece of information for any tracking study, as they contain important information about each data file, such as tag ID, animal ID, or deployment information, that we can add back into to our raw data when needed. For example, the table below shows what the first few columns of the metadata file looks like for our example red-footed booby data:

| TagID | BirdID | DeployID | Species | Population | Age | BreedingStage | DeploymentDate | DeploymentTime | RetrievalDate |
|---|---|---|---|---|---|---|---|---|---|
| 5 | GV37501 | 1 | RFB | DG | Adult | Chick rearing | 25/06/2016 | 16:25:00 | 29/06/2016 |
| 46 | GV37503 | 1 | RFB | DG | Adult | Chick rearing | 26/06/2016 | 08:30:00 | 01/07/2016 |
| 35 | GV37734 | 1 | RFB | NI | Adult | Chick rearing | 08/07/2018 | 09:48:00 | 11/07/2018 |

#### Select file and date/time formats

🧠 **User input required**

First we define the path to our metadata file:

```
filepath_meta <- here("Data","RFB_Metadata.csv")
```

Then much like in Step 1, we define the date format(s) used (for passing to `lubridate`) (`d` = day as decimal, `m` = month as decimal, `y` = year without century - 2 digits, `Y` = year with century - 4 digits). Here, we've included common combinations, which you'll need to modify if your metadata include a different format (run `OlsonNames()` to return a full list of time zones names).

```
metadate_formats <- c("dmY", "Ymd") #specify date format used in metadata
metadatetime_formats <- c("dmY HMS", "Ymd HMS") #specify date & time format
metadatatimezone <- "Indian/Chagos" #specify timezone used for metadata
```

Next we read in the metadata file (make sure to check the `read_` function you're using matches your data format!).

```
df_metadata <- readr::read_csv(filepath_meta) # Read in metadata file
names(df_metadata)
```

#### Select metadata columns

🧠 **User input required**

Then we select necessary columns & create a new complete dataframe, making sure to provide four compulsory columns: **ID** — as defined in tracking data (individual **ID** or **TagID**), **deployment date** & **deployment time**. We can also provide optional columns depending on sensor type: e.g. colony, sex, age. You can add or delete other columns where appropriate.

If you have multiple ID columns like TagID/DeployID, include them here (for example, if one individual was tracked over multiple deployments/years, or if one tag was re-deployed on multiple individuals). For more information and helpful code, see the FAQ document and troubleshooting script.

**Deployment and retrieval dates:** Different tags types sometimes require specific approaches for dealing with data collected outside of deployment period (e.g., before deployment or after retrieval). If data need to be filtered for one or both of these scenarios, we need to sort out these columns in the metadata, and if not relevant for the data, set the column name to "NA".

**Central Place foragers:** If you are working with a central place forager (e.g., animals returning to a breeding location) and you have individual breeding location information in your metadata, here is a good place to add this info to the tracking data (e.g., breeding seabirds with known individual nest location, or seals returning to known haul-out location). We recommend adding these columns as: `CPY = Central place Y coordinate column` & `CPX = Central place X coordinate column`

```
df_metadataslim <- data.frame(ID = as.character(df_metadata$BirdID), # compulsory column
                              TagID = as.character(df_metadata$TagID),
                              DeployID = as.character(df_metadata$DeployID),
                              DeployDate_local = df_metadata$DeploymentDate, # compulsory colu
                              DeployTime_local = df_metadata$DeploymentTime, # compulsory colu
                              RetrieveDate_local = df_metadata$RetrievalDate, # compulsory co
                              RetrieveTime_local = df_metadata$RetrievalTime, # compulsory co
                              CPY = df_metadata$NestLat,
                              CPX = df_metadata$NestLong,
                              Species = "RFB",
                              Population = df_metadata$Population,
                              Age = df_metadata$Age,
                              BreedStage = df_metadata$BreedingStage)
```

**Option: select alternative columns**

Format all dates and times, combine them and specify timezone (NA's in deployment/retrieval date times will throw warnings, but these are safe to ignore if you know there are NA's in these columns).

```
df_metadataslim <- df_metadataslim %>%
  mutate(Deploydatetime =
           lubridate::parse_date_time(
             paste(DeployDate_local, DeployTime_local),# make deploy datetime
                                          order = metadatetime_formats,
                                          tz = metadatatimezone),
         Retrievedatetime =
           lubridate::parse_date_time(
             paste(RetrieveDate_local, RetrieveTime_local), # make retrieve datetime
                                            order=metadatetime_formats,
                                            tz=metadatatimezone)
         ) %>%
  select(-any_of(c("DeployDate_local",
                   "DeployTime_local",
                   "RetrieveDate_local",
                   "RetrieveTime_local"))
         ) %>%
  mutate(across(contains('datetime'), # for chosen datetime column
                ~with_tz(., tzone = trackingdatatimezone)) #format to different tz
         )
```

Here we'll create a dataframe of temporal extents of our data to use in absence of deploy/retrieve times (this is also useful for basic data checks and for writing up methods).

```
df_temporalextents <- df_raw %>%
  group_by(across(all_of(ID_type))) %>%
  summarise(min_datetime = min(DateTime),
            max_datetime = max(DateTime))
```

Then we use these temporal extents of our data to fill in any NA's in the deploy/retrieve times.

```
df_metadataslim <- df_metadataslim %>%
  left_join(., df_temporalextents, by = ID_type) %>%
```

```
  mutate(Deploydatetime = case_when(!is.na(Deploydatetime) ~ Deploydatetime,
                                    is.na(Deploydatetime) ~ min_datetime),
         Retrievedatetime = case_when(!is.na(Retrievedatetime) ~ Retrievedatetime,
                                    is.na(Retrievedatetime) ~ max_datetime)) %>%
  select(-c(min_datetime, max_datetime))
```

Next we merge metadata with raw data using the ID column.

```
df_metamerged <- df_raw %>%
  left_join(., df_metadataslim, by=ID_type)
```

Finally, we can remove intermediate files/objects by specifying objects to keep.

```
rm(list=ls()[!ls() %in% c("df_metamerged", "species_code")]) #specify objects to keep
```

### 3. Cleaning

🧠 **User input required**

Define your own no/empty/erroneous data values in `Lat` and `Lon` columns (e.g. “bad” values specified by the tag manufacturer).

```
No_data_vals <- c(0, -999)
```

Define a vector of columns which can’t have `NA` s (if there are `NA` s in one of these columns the problematic row will be removed).

```
na_cols <- c("X", "Y", "DateTime", "ID")
```

Now we pipe the data through a series of functions to drop `NA` s in specified columns, filter out user-defined `no_data_values` in `Lat Lon` columns, remove duplicates, remove undeployed locations and filter out locations within temporal cut-off following deployment.

```
df_clean <- df_metamerged %>%
  drop_na(all_of(na_cols)) %>%
  filter(!X %in% No_data_vals & !Y %in% No_data_vals) %>% # remove bad Lat/Lon values
  distinct(DateTime, ID, .keep_all = TRUE) %>% # NB: might be an issue for ACC without
  filter(
    case_when(!is.na(Retrievedatetime) # for all valid datetimes
              ~ Deploydatetime < DateTime & # keep if datetime after deployment...
                DateTime < Retrievedatetime, # ...and before retrieval
              .default = Deploydatetime < DateTime)) # filter deployment only if retri
head(df_clean)
```

```
       ID       Date     Time         Y        X            DateTime Year
1 GV37501 2016-06-25 10:27:56 -7.238936 72.435043 2016-06-25 10:27:56 2016
2 GV37501 2016-06-25 10:33:13 -7.238903 72.435005 2016-06-25 10:33:13 2016
3 GV37501 2016-06-25 10:38:19 -7.238754 72.434944 2016-06-25 10:38:19 2016
4 GV37501 2016-06-25 10:43:46 -7.238957 72.435188 2016-06-25 10:43:46 2016
5 GV37501 2016-06-25 10:49:02 -7.239048 72.435059 2016-06-25 10:49:02 2016
```

```
6 GV37501 2016-06-25 10:54:38 -7.238441 72.434708 2016-06-25 10:54:38 2016
  TagID DeployID      CPY      CPX Species Population   Age    BreedStage
1     5        1 -7.2386 72.4347     RFB         DG Adult Chick rearing
2     5        1 -7.2386 72.4347     RFB         DG Adult Chick rearing
3     5        1 -7.2386 72.4347     RFB         DG Adult Chick rearing
4     5        1 -7.2386 72.4347     RFB         DG Adult Chick rearing
5     5        1 -7.2386 72.4347     RFB         DG Adult Chick rearing
6     5        1 -7.2386 72.4347     RFB         DG Adult Chick rearing
       Deploydatetime    Retrievedatetime
1 2016-06-25 10:25:00 2016-06-29 03:32:00
2 2016-06-25 10:25:00 2016-06-29 03:32:00
3 2016-06-25 10:25:00 2016-06-29 03:32:00
4 2016-06-25 10:25:00 2016-06-29 03:32:00
5 2016-06-25 10:25:00 2016-06-29 03:32:00
6 2016-06-25 10:25:00 2016-06-29 03:32:00
```

**Option: Filter by fix quality**

Finally we remove intermediate files/objects:

```
rm(list=ls()[!ls() %in% c("df_clean", "species_code")]) #specify objects to keep
```

### 4. Processing

#### Perform some useful temporal and spatial calculations on the data

🧠 **User input required**

First we need to specify the co-ordinate projection systems for the tracking data and meta data. The default here is lon/lat for both tracking data & metadata, for which the EPSG code is 4326. For more information see the CRS section of the FAQ's or have a look at the ESPG.io database.

```
tracking_crs <- 4326 # Only change if data are in a different coordinate system
meta_crs <- 4326 # Only change if data are in a different coordinate system
```

Next we transform coordinates of data, and perform spatial calculations. This requires spatial analysis, and so it is good practice to run all spatial analyses in a coordinate reference system that uses metres as a unit.

The default CRS for this workflow is the Spherical Mercator projection — (`crs = 3857`), which is used by Google Maps/OpenStreetMap and works worldwide. However, this CRS can over-estimate distance calculations in some cases, so it's important to consider the location and scale of your data (e.g., equatorial/polar/local scale/global scale) and choose a projection system to match. Other options include (but are not limited to) UTM, and Lambert Azimuthal Equal-Area (LAEA) projections.

```
transform_crs <- 3857
```

Here we'll calculate some useful movement metrics from the tracking data, including distance between fixes, time between fixes, and net displacement from the first fix.

```
df_diagnostic <-  df_clean %>%
  ungroup() %>% #need to ungroup to extract geometry of the whole dataset
  mutate(geometry_GPS = st_transform( # transform X/Y coordinates
            st_as_sf(., coords=c("X","Y"), crs = tracking_crs), #from original format
            crs = transform_crs)$geometry # to the new transform_crs format
         ) %>%
  group_by(ID) %>% #back to grouping by ID for calculations per individual
  mutate(dist = st_distance(geometry_GPS, # distance travelled from previous fix,
                            lag(geometry_GPS),
                            by_element = T), # calculations are done by row
         difftime = difftime(DateTime, lag(DateTime), # time passed since previous fix
                             units = "secs"), # in seconds
         netdisp = st_distance(geometry_GPS, # dist. between 1st and current location
                               geometry_GPS[1],
                               by_element = F)[,1], # dense matrix w/ pairwise distanc
         speed = as.numeric(dist)/as.numeric(difftime), # calculate speed (distance/ti
         dX = as.numeric(X)-lag(as.numeric(X)), #diff. in lon  relative to prev. locat
         dY = as.numeric(Y)-lag(as.numeric(Y)), #diff. in lat relative to prev. locati
         turnangle = atan2(dX, dY)*180/pi + (dX < 0)*360) %>% # angle from prev. to cu
  ungroup() %>%
  select(-c(geometry_GPS, dX, dY)) # ungroup and remove excess geometries
```

Add latitude and longitude column — this can be useful for plotting and is a common coordinate system used in the shiny app

```
df_diagnostic <- st_coordinates(st_transform(st_as_sf(df_diagnostic,
                                                      coords = c("X","Y"),
                                                      crs=tracking_crs),
                                             crs = 4326)) %>%
                 as.data.frame() %>%
                 rename("Lon" = "X", "Lat" = "Y") %>%
                 cbind(df_diagnostic, .)
```

#### 5. Save for Shiny

Here we're going to save `df_diagnostic` to use in the Shiny app provided. The app is designed to explore how further filtering and processing steps affect the data.

🧠 **User input required**

First, we use here to create a file path for saving the working dataframe files, and create the folder if missing

```
filepath_dfout <- here("DataOutputs","WorkingDataFrames") # create filepath
dir.create(filepath_dfout) # create folder if it doesn't exist
```

Next we define file name for the saved file by pasting the species code before `_diagnostic` (can change this if you want to use a different naming system).

```
filename_dfout <- paste0(species_code, "_diagnostic")
```

> If not added from the metadata, add a species column and any other columns here relevant to your data *(optional)*
>
> ```
> ## ** Option ** ##
> df_diagnostic$Species <- species_code
> ```

Finally we save the `df_diagnostic` as a csv file using the variables created above.

```
write_csv(df_diagnostic, file = here(filepath_dfout, paste0(filename_dfout,".csv")))
```

Remove everything except `df_diagnostic` ahead of the next step.

```
rm(list=ls()[!ls() %in% c("df_diagnostic", "species_code")]) #specify objects to keep
```

### 6. Filtering

This second filtering stage is designed to remove outliers in the data, and you can use outputs from the Shiny app to inform these choices. If you don't need to filter for outliers, skip this step and keep using `df_diagnostic` in the next steps.

#### Accessing the Shiny app

##### Option 1:

Access the Shiny app online at the following link: [https://lukeozsanlav.shinyapps.io/exmove_explorer/](https://lukeozsanlav.shinyapps.io/exmove_explorer/)

##### Option 2:

Alternatively run the app from your local R session with the following code

```
if (!require("shiny")) install.packages("shiny")
library(shiny)
runGitHub("ExMoveApp", username = "LukeOzsanlav",
          ref = "master", subdir = "app")
```

##### App usage:

- Upload your csv version of `df_diagnostic` to the app by clicking the `Upload data` button in the top left.
- At the bottom of each app page are printed code chunks that can be copied into subsequent user input section. These code chunks contain the user input values you manually select in the app

##### Define threshold values

For this section, you can either use the code chunks produced by the Shiny app, or manually define the threshold values yourself. If you're using the Shiny app, you can copy the code chunks from the bottom of each page to replace the user input section below. If you're manually defining the threshold values, you can edit each of the variables below as you would for any of the user input sections (we have suggested some for the RFB dataset).

🧠 **User input required**

First we define a period to filter after tag deployment, when all points before the cutoff will be removed (e.g. to remove potentially unnatural behaviour following the tagging event). We define this period using the `as.period` function, by providing an integer value and time unit (e.g. hours/days/years). This code below specifies a period of 30 minutes:

```
filter_cutoff <- as.period(30, unit="minutes")
```

Then we define speed threshold in m/s, which we will use to remove any points with faster speeds.

```
filter_speed <- 20
```

Next we define a net displacement (distance from first point) threshold and specify units. Any points further away from the first tracking point will be removed (see commented code for how to retain all points):

```
filter_netdisp_dist <- 300
filter_netdist_units <- "km" # e.g., "m", "km"

#If you want to retain points no matter the net displacement value, use these values instead:
#filter_netdisp_dist <- max(df_diagnostic$netdisp)
#filter_netdist_units <- "m"
```

##### Implement filters

Create net displacement filter using distance and units

```
filter_netdisp <- units::as_units(filter_netdisp_dist, filter_netdist_units)
```

Filter df_diagnostic

```
df_filtered <- df_diagnostic %>%
  filter(Deploydatetime + filter_cutoff < DateTime, # keep times after cutoff
         speed < filter_speed, # keep speeds slower than speed filter
         netdisp <= filter_netdisp) # keep distances less than net displacement filter
head(df_filtered)
```

```
       ID       Date     Time         Y        X            DateTime Year
1 GV37501 2016-06-25 11:01:04 -7.238462 72.434708 2016-06-25 11:01:04 2016
2 GV37501 2016-06-25 11:08:50 -7.238486 72.434738 2016-06-25 11:08:50 2016
3 GV37501 2016-06-25 11:17:13 -7.238597 72.435005 2016-06-25 11:17:13 2016
4 GV37501 2016-06-25 11:25:16 -7.231193 72.434624 2016-06-25 11:25:16 2016
5 GV37501 2016-06-25 11:30:32  -7.23855 72.434731 2016-06-25 11:30:32 2016
6 GV37501 2016-06-25 11:36:43 -7.238533 72.434761 2016-06-25 11:36:43 2016
  TagID DeployID     CPY      CPX Species Population   Age     BreedStage
1     5        1 -7.2386 72.4347     RFB          DG Adult Chick rearing
2     5        1 -7.2386 72.4347     RFB          DG Adult Chick rearing
3     5        1 -7.2386 72.4347     RFB          DG Adult Chick rearing
4     5        1 -7.2386 72.4347     RFB          DG Adult Chick rearing
```

```
5     5        1 -7.2386 72.4347     RFB         DG Adult Chick rearing
6     5        1 -7.2386 72.4347     RFB         DG Adult Chick rearing
       Deploydatetime     Retrievedatetime          dist difftime       netdisp
1 2016-06-25 10:25:00 2016-06-29 03:32:00   2.356490 [m] 386 secs  64.96002 [m]
2 2016-06-25 10:25:00 2016-06-29 03:32:00   4.290196 [m] 466 secs  60.84931 [m]
3 2016-06-25 10:25:00 2016-06-29 03:32:00  32.226707 [m] 503 secs  38.27498 [m]
4 2016-06-25 10:25:00 2016-06-29 03:32:00 831.906239 [m] 483 secs 870.11594 [m]
5 2016-06-25 10:25:00 2016-06-29 03:32:00 825.636254 [m] 316 secs  55.51973 [m]
6 2016-06-25 10:25:00 2016-06-29 03:32:00   3.846024 [m] 371 secs  55.05007 [m]
        speed turnangle      Lon       Lat
1 0.006104896 180.00000 72.43471 -7.238462
2 0.009206429 128.65981 72.43474 -7.238486
3 0.064069000 112.57414 72.43501 -7.238597
4 1.722373165 357.05423 72.43462 -7.231193
5 2.612772957 179.16675 72.43473 -7.238550
6 0.010366640  60.46122 72.43476 -7.238533
```

Remove intermediate files/objects

```
rm(list=ls()[!ls() %in% c("df_filtered", "species_code")]) #specify objects to keep
```

### 7. Summarise cleaned & filtered tracking data

🧠 **User input required**

Set the units to display sampling rate in the summary table

```
sampleRateUnits <- "mins"
```

#### Define levels of grouping factors to summarise over

Firstly, down to population level. Here, we are working on data from one population & year, and so use `Species` as the grouping factor. Add any other relevant grouping factors here (e.g. Country / Year / Season / Age).

```
grouping_factors_poplevel <- c("Species")
```

Secondly, down to individual level (add `DeployID` for example if relevant).

```
grouping_factors_indlevel <- c("ID")
```

#### Create summary tables

Create a small function to calculate standard error.

```
se <- function(x) sqrt(var(x, na.rm = T) / length(x[!is.na(x)]))
```

Create a summary table of individual-level summary statistics:

```
df_summary_ind <- df_filtered %>%
  group_by(across(c(all_of(grouping_factors_poplevel), all_of(grouping_factors_indleve
  summarise(NoPoints = NROW(ID), # number of fixes
```

```
NoUniqueDates = length(unique(Date)), # number of tracking dates
          FirstDate = as.Date(min(Date)), # first tracking date
          LastDate = as.Date(max(Date)), # last tracking date
          SampleRate = mean(as.numeric(difftime, units = sampleRateUnits), na.rm = T
          SampleRate_se = se(as.numeric(difftime, units = sampleRateUnits))) # sampl
```

```
`summarise()` has grouped output by 'Species'. You can override using the
`.groups` argument.
```

```
df_summary_ind
```

```
# A tibble: 3 × 8
# Groups:   Species [1]
  Species ID      NoPoints NoUniqueDates FirstDate  LastDate   SampleRate
  <chr>   <chr>      <int>         <int> <date>     <date>          <dbl>
1 RFB     GV37501      989             5 2016-06-25 2016-06-29       5.37
2 RFB     GV37503     1280             6 2016-06-26 2016-07-01       5.58
3 RFB     GV37734      820             4 2018-07-08 2018-07-11       5.18
# ℹ 1 more variable: SampleRate_se <dbl>
```

Create a table of population-level summary statistics:

```
df_summary_pop <- df_summary_ind %>% # use the individual-level summary data
  group_by(across(grouping_factors_poplevel)) %>%
  summarise(NoInds = length(unique(ID)), # number of unique individuals
            NoPoints_total = sum(NoPoints), # total number of tracking locations
            FirstDate = as.Date(min(FirstDate)), # first tracking date
            LastDate = as.Date(max(LastDate)), # last tracking date
            PointsPerBird = mean(NoPoints), # number of locations per individual: mean
            PointsPerBird_se = se(NoPoints), # number of locations per individual: sta
            DatesPerBird = mean(NoUniqueDates), # number of tracking days per bird: me
            DatesPerBird_se = se(NoUniqueDates), # number of tracking days per bird: s
            SampleRate_mean = mean(SampleRate), # sample rate mean
            SampleRate_se = se(SampleRate)) # sample rate standard error
df_summary_pop
```

```
# A tibble: 1 × 11
  Species NoInds NoPoints_total FirstDate  LastDate   PointsPerBird
  <chr>    <int>          <int> <date>     <date>             <dbl>
1 RFB          3           3089 2016-06-25 2018-07-11         1030.
# ℹ 5 more variables: PointsPerBird_se <dbl>, DatesPerBird <dbl>,
#   DatesPerBird_se <dbl>, SampleRate_mean <dbl>, SampleRate_se <dbl>
```

Remove intermediate files/objects by specifying which objects to keep:

```
rm(list=ls()[!ls() %in% c("df_filtered",
                          "df_summary_ind", "df_summary_pop",
                          "species_code")])
```

#### 8. Save filtered and summary data

🧠 User input required

First we define the folder file path for saving our filtered data and create folder if not already present

```
filepath_filtered_out <- here("DataOutputs","WorkingDataFrames")
dir.create(filepath_filtered_out)
```

Then we define the file path for saving summary dataframes, again creating folder if needed

```
filepath_summary_out <- here("DataOutputs","SummaryDataFrames")
dir.create(filepath_summary_out)
```

Here we define file names for saved files, and paste the species code to `_summary_`, followed by `ind` (individual level) or `pop` (population level). You can change this if you want to use a different naming system.

```
filename_filtered_out <- paste0(species_code, "_filtered")
filename_summary_ind_out <- paste0(species_code, "_summary_ind")
filename_summary_pop_out <- paste0(species_code, "_summary_pop")
```

Now we can save all our dataframes as .csv files using our defined values

```
write_csv(df_filtered, file = here(filepath_filtered_out, paste0(filename_filtered_out
write_csv(df_summary_ind, file = here(filepath_summary_out, paste0(filename_summary_in
write_csv(df_summary_pop, file = here(filepath_summary_out, paste0(filename_summary_po
```

Lastly we remove intermediate files/objects

```
rm(list=ls()[!ls() %in% c("df_filtered",
                          "df_summary_ind", "df_summary_pop",
                          "species_code")])
```

### 9. Visualisation

User input required

Define parameters for reading out plots, and define device to read plots out as e.g. tiff/jpeg

```
device <- "tiff"
```

Define units for plot size (usually mm)

```
units <- "mm"
```

Define plot resolution in dpi (300 is usually good minimum)

```
dpi <- 300
```

Define filepath to read out plots and create folder if absent

```
out_path <- here("DataOutputs","Figures")
dir.create(out_path)
```

We plot maps over a topography base-layer which can include terrestrial (elevation) and marine (bathymetry/water depth) data. To set legend label for topography data, relevant to your data.

```
topo_label = "Depth (m)"
```

Load additional libraries for spatial visualisation **(optional)**

**If you see a `masking` warning these are fine. Watch out for packages that aren't installed yet**

```
library(rnaturalearth)
library(marmap)
library(plotly)
```

Create version of data for plotting by transforming required columns to numeric and creating time elapsed columns

```
df_plotting <- df_filtered %>%
  group_by(ID) %>%
  mutate(diffsecs = as.numeric(difftime),
         secs_elapsed = cumsum(replace_na(diffsecs, 0)),
         time_elapsed = as.duration(secs_elapsed),
         days_elapsed = as.numeric(time_elapsed, "days")) %>%
  mutate(across(c(dist,speed, Lat, Lon), as.numeric))
```

Create a map of all points. Set the plot limits as the max and min lat/longs as the tracking data

First set up a basemap to plot over: - Use `rnaturalearth` low resolution countries basemap - co-ordinates in lat/lon to match other spatial data

```
countries <- ne_countries(scale = "medium", returnclass = "sf")
```

Define min and max co-ordinates based on extent of tracking data, for adding bathymetry extracted from NOAA database.

```
minlon <- min(df_plotting$Lon)
maxlon <- max(df_plotting$Lon)

minlat <- min(df_plotting$Lat)
maxlat <- max(df_plotting$Lat)
```

Load in bathymetry basemap. Set limits slightly beyond tracking data to make a buffer so no gaps when plotting

```
base_topography_map <- getNOAA.bathy(
  lon1 = minlon - 0.1, lon2 = maxlon + 0.1,
```

```
  lat1 = minlat - 0.1, lat2 = maxlat + 0.1,
  resolution = 1)
```

Fortify bathymetry data for plotting

```
base_topography_fort = fortify(base_topography_map)
```

Create base map with correct extent, topography, country outlines, etc.,

```
map_base <- ggplot() +
  geom_raster(data = base_topography_fort, aes(x=x, y=y, fill=z), alpha = 0.9) +
  # add colour scheme for the fill
  scale_fill_viridis_c(option="mako", name = topo_label) +
  # add map of countries over the top
  geom_sf(data = countries, aes(geometry = geometry), fill = NA) +
  # set plot limits
  coord_sf(xlim = c(minlon-0.1, maxlon+0.1),
           ylim = c(minlat-0.1, maxlat+0.1), crs = 4326, expand = F) +
  # add labels
  labs(x = "Longitude", y = "Latitude") +
  theme(axis.text=element_text(colour="black"),
        axis.title.x = element_text(size = 15),
        axis.text.x = element_text(hjust=0.7),
        axis.title.y = element_text(angle=90, vjust = 0.4, size = 15),
        axis.text.y = element_text(hjust=0.7, angle=90, vjust=0.3)) +
  # set a theme
  theme_light()
map_base
```

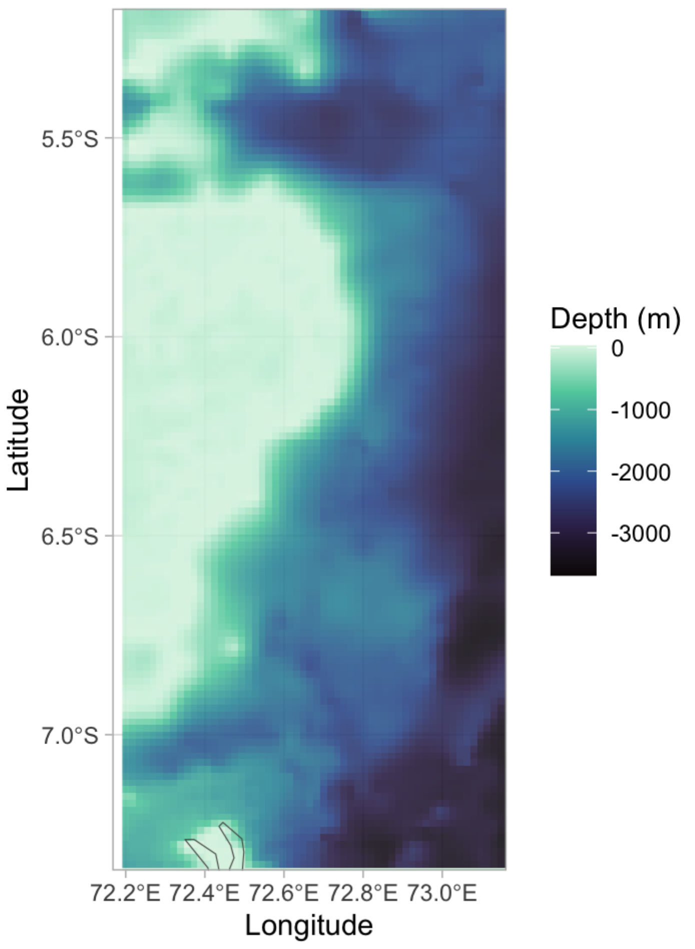

Population | Several individuals | Option: Many individuals

Plot a combined map of all tracking locations:

```
map_alllocs <- map_base +
  # add GPS points
  geom_point(data = df_plotting,
             aes(x = Lon, y = Lat),
             alpha = 0.8, size = 0.5, col = "violetred3")
map_alllocs
```

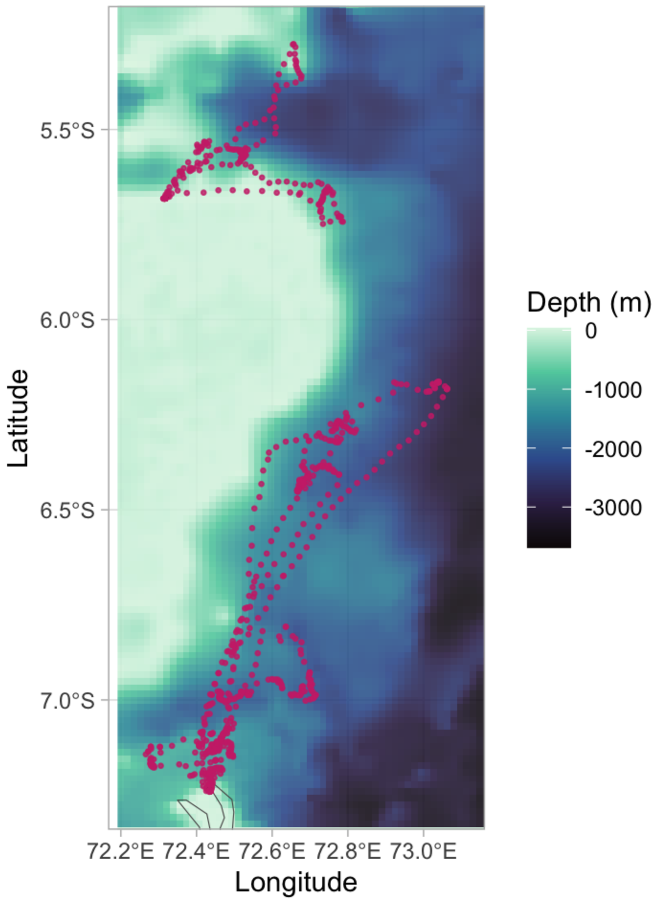


Save maps for further use using `ggsave` function.

```
ggsave(plot = map_alllocs,
       filename = paste0(species_code, "_map_all_locs.", device),
       device = device,
       path = out_path,
       units = units, width = 200, height = 175, dpi = dpi,
)

ggsave(plot = map_individuals,
       filename = paste0(species_code, "_map_individuals.", device),
       device = device,
       path = out_path,
```

```
        units = units, width = 200, height = 175, dpi = dpi,
)
```

Create a time series plot of speed, faceted for each individual.

```
speed_time_plot <- df_plotting %>% #speed over time
  ggplot(data = .,
         aes(x = days_elapsed, y = speed, group = ID)
         ) +
  geom_line() +   # add line of speed over time
  xlab("time elapsed (days)") +
  ylab("speed (m/s)") +
  facet_wrap(~ID, # facet by individual
             nrow = round(sqrt(n_distinct(df_plotting$ID)))) +
  theme_light() + # set plotting theme
  theme(axis.text = element_text(colour="black")) #adjust theme
speed_time_plot
```

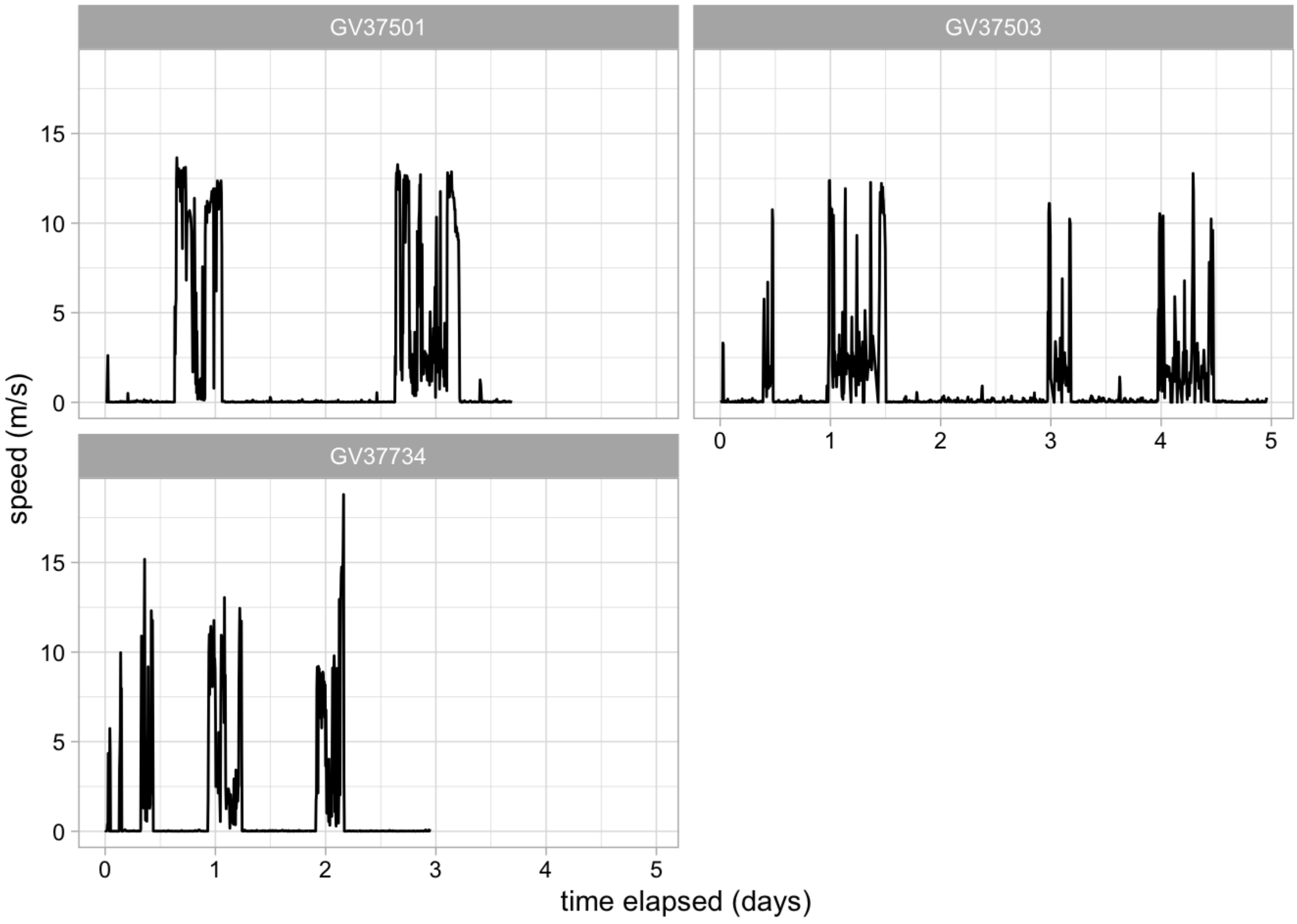


> **Note**
>
> Warnings about `non-finite` values for speed/step length plots are expected and usually refer to the first location for each individual (i.e. number of non-finite values should be equal to number of individuals)

Save plot for further use

```
ggsave(plot = speed_time_plot,
       filename = paste0(species_code, "_speed_timeseries_plot.", device),
```

```
        device = device,
        path = out_path,
        units = units, width = 200, height = 175, dpi = dpi
)
```

Create a histogram of point to point speeds (can adjust binwidth and x limits manually)

```
speed_hist <- df_plotting %>%
  ggplot(data = ., aes(speed)) +
  geom_histogram(binwidth = 0.1, alpha=0.7) + # can adjust binwidth to suite your need
  geom_density(aes(y = 0.1*..count..)) +
  xlab("speed (m/s)") +
  ylab("count") +
  facet_wrap(~ID, # facet by individual
             nrow = round(sqrt(n_distinct(df_plotting$ID)))) +
  theme_light() + # set plotting theme
  theme(axis.text = element_text(colour="black")) #adjust theme
speed_hist
```

```
Warning: The dot-dot notation (`..count..`) was deprecated in ggplot2 3.4.0.
ℹ Please use `after_stat(count)` instead.
```

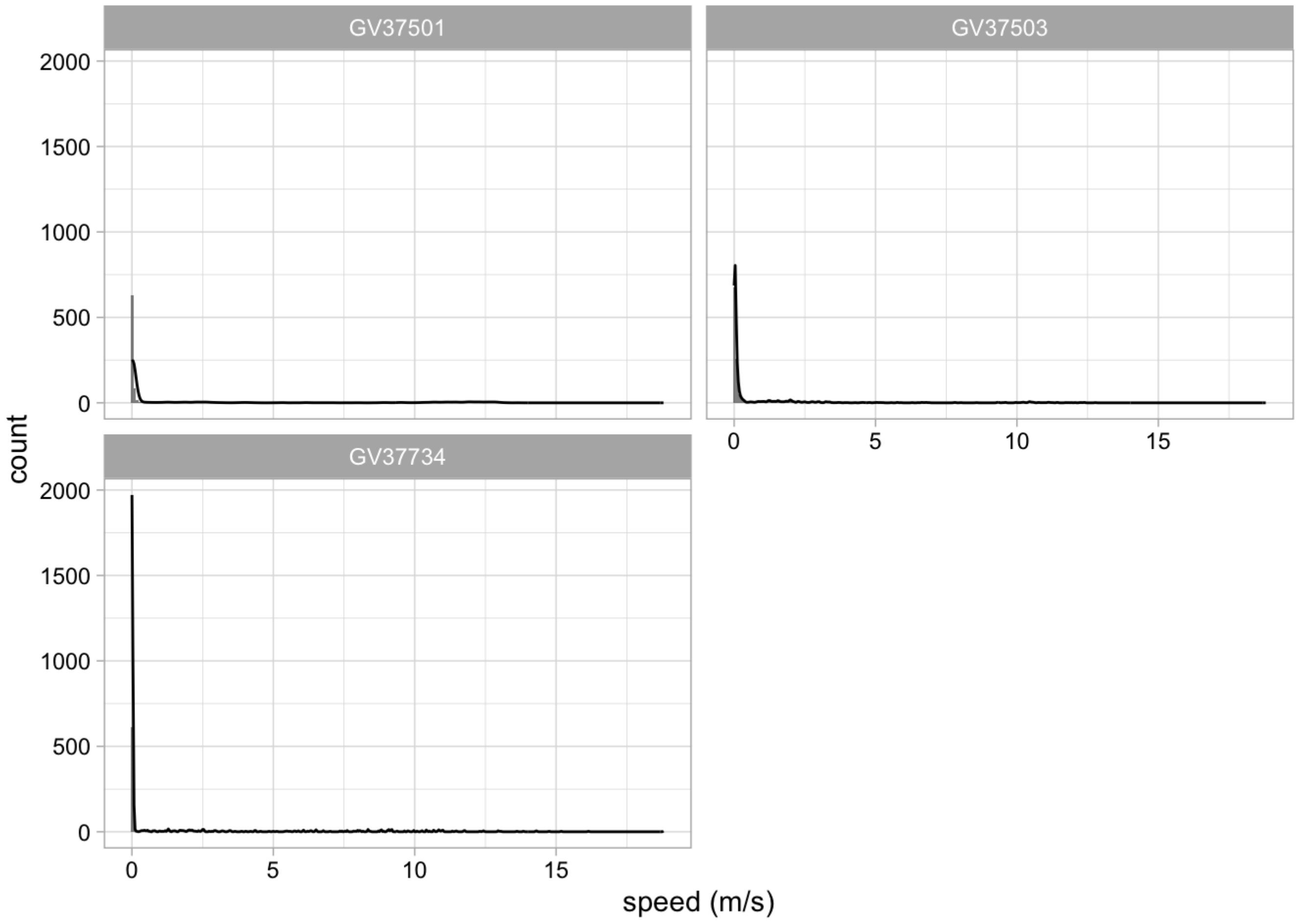


Save plot for further use

```
ggsave(plot = speed_hist,
        filename = paste0(species_code, "_speed_histogram", device),
        device = device,
        path = out_path,
```

```
        units = units, width = 200, height = 175, dpi = dpi,
)
```

Create a time series plot of step lengths (faceted for each individual)

```
step_time_plot <- df_plotting %>% #step length over time
  ggplot(data = .,
         aes(x = days_elapsed, y = as.numeric(netdisp), group = ID)) +
  geom_line() +
  # add plot labels
  xlab("time elapsed (days)") + ylab("Distance from first fix (m)") +

  facet_wrap(~ID, # facet by individual
             nrow= round(sqrt(n_distinct(df_plotting$ID)))) +
  theme_light() + # set plotting theme
  theme(axis.text = element_text(colour="black")) #adjust theme
step_time_plot
```

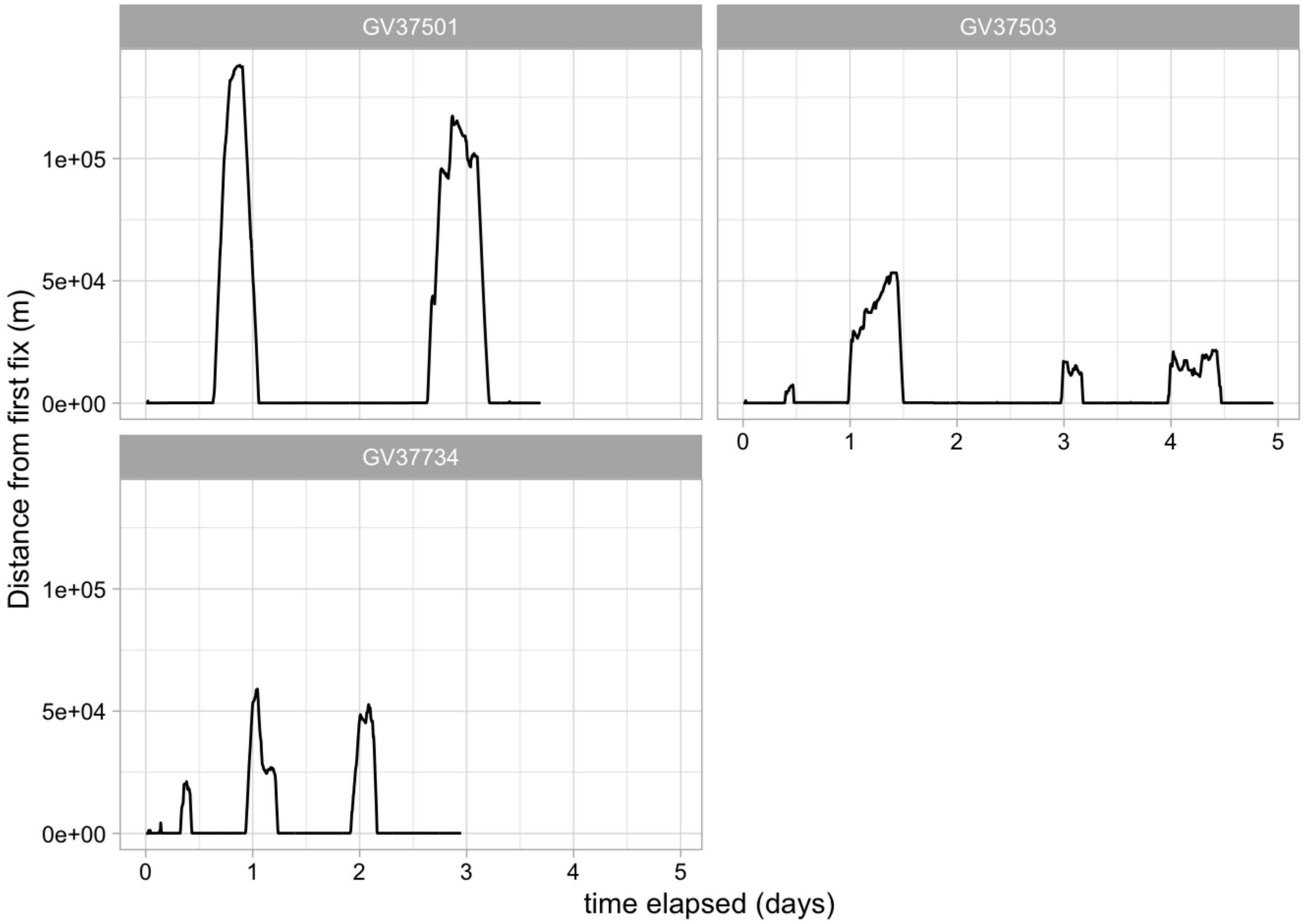


Save plot for further use

```
ggsave(plot = step_time_plot,
       filename = paste0(species_code, "_step_time_plot.", device),
       device = device,
       path = out_path,
       units = units, width = 200, height = 175, dpi = dpi,
)
```

Create a histogram of step lengths (can adjust binwidth and x limits manually)

```
step_hist <- df_plotting %>% #step histogram
  ggplot(data = .,
         aes(as.numeric(dist))) +
  geom_histogram(binwidth = 1, alpha = 0.7) + # can adjust binwidth to suite your need
  geom_density(aes(y = 1*..count..)) +
  xlab("step length (m)") +
  ylab("count") +
  facet_wrap(~ID, # facet by individual
             nrow = round(sqrt(n_distinct(df_plotting$ID))))+
  theme_light() # set plotting theme
```

Save plot for further use

```
ggsave(plot = step_hist,
       filename = paste0(species_code, "_step_hist.", device),
       device = device,
       path = out_path,
       units = units, width = 200, height = 175, dpi = dpi,
)
```

Lastly, we remove intermediate files/objects if necessary to speed up any post-processing steps

```
rm(list=ls()[!ls() %in% c("species_code")]) #specify objects to keep
```
